# Bone elongation in the embryo occurs without column formation in the growth plate

**DOI:** 10.1101/2023.11.14.567062

**Authors:** Sarah Rubin, Ankit Agrawal, Anne Seewald, Paul Villoutreix, Adrian Baule, Elazar Zelzer

## Abstract

Chondrocyte columns, which are a hallmark of growth plate architecture, play a central role in bone elongation. Columns are formed by clonal expansion following rotation of the division plane, resulting in a stack of cells oriented parallel to the growth direction. However, despite decades of research, column structure has thus far been studied only in two dimensions. To fill this knowledge gap, we analyzed hundreds of Confetti multicolor clones in growth plates of mouse embryos using a pipeline comprising 3D imaging and algorithms for morphometric analysis. Surprisingly, analysis of the elevation angles between neighboring pairs of cells revealed that most cells did not display the typical stacking pattern associated with column formation, implying incomplete rotation of the division plane. Morphological analysis revealed that although embryonic clones were elongated, they formed clusters oriented perpendicular to the growth direction. Analysis of growth plates of postnatal mice revealed both complex columns, composed of both ordered and disordered cell stacks, and small, disorganized clusters located in the outer edges. Our finding that embryonic growth plates function without forming columns suggests that longitudinal bone growth is regulated by different cellular mechanisms during pre- and postnatal development. Moreover, the observed complex columnar and cluster arrangements may serve other, yet unknown morphogenetic functions. More broadly, our findings provide a new understanding of the cellular mechanisms underlying growth plate activity and bone elongation during development.

## INTRODUCTION

Cellular organization plays a major role in tissue and organ morphogenesis (*1–6*). The mammalian growth plate is an excellent example for this concept, as its complex architecture is the engine driving longitudinal bone growth (*7–11*). The growth plate, which is located at both ends of developing long bones, drives bone elongation by a tightly regulated process of cell proliferation and differentiation, which involves increase in cell size and their organization along the proximal-distal (P-D) axis (*12–15*). The growth plate comprises four zones. Most extreme is the resting zone (RZ), where chondrocytes are small and disorganized. Underneath lies the proliferative zone (PZ), where chondrocytes increase in volume, adopt a flat and elongated morphology, and organize into columns (*16–20*). In the subsequent prehypertrophic (PHZ) and hypertrophic zones (HZ), cells reach their maximum size (*7, 11, 21*). These changes in cell size and spatial organization determine the rate of bone elongation (*8–10, 22–24*).

Columnar arrangement of chondrocytes has been a subject of study for nearly a century (*17*), gaining attention due to the remarkable emergence of cellular order from the highly disordered RZ. This columnar arrangement facilitates bone elongation by maximizing cell density in the longitudinal axis while limiting it laterally, thereby constraining hypertrophic cell growth to the P-D axis (*25*). In the PZ, the division of column-forming cells is perpendicular to the P-D axis. Considering that these cells ultimately orient themselves with their short axis parallel to the P-D axis, the rearrangement into elongated columns requires a robust morphogenetic mechanism. Originally, analyses of two-dimensional static images suggested that in the embryonic growth plate, columns form through a process akin to convergent extension, an evolutionarily conserved tissue elongation mechanism involving cell intercalation (*18, 26–29*). However, more recent live imaging studies in various model systems showed that cells do not intercalate to form columns (*19, 25, 30*). Instead, following cell division, sister cells undergo a cell-cell and cell-extracellular matrix (ECM) adhesion-dependent 90° rotation prior to separation. This rotation ensures that cells are neatly stacked with their short axis parallel to the P-D axis.

Recent studies have highlighted three fundamental principles governing column formation. First, columns consist of clonal cells (*18–20, 26, 31–33*). Whereas embryonic columns are multiclonal, postnatally, following the formation of secondary ossification centers, columns become monoclonal and originate from Pthrp+ RZ cells (*31–33*). The second principle is that cells within the column orient their short axis parallel to the P-D axis of the bone (*7, 18–20, 26, 27, 34*) within a threshold of 12° (*19*). The third rule pertains to the alignment of the column itself. The long axis is oriented parallel to the P-D axis (*17, 18, 20, 26, 27, 34, 35*) within a 12° threshold for single columns and a 20° threshold for complex columns (*19*).

Over the years, numerous studies have been dedicated to deciphering the molecular and cellular processes underpinning the formation of columns and their involvement in bone elongation. Studies in embryonic and postnatal mouse limbs have shown the importance of interactions between chondrocytes and the surrounding ECM. These studies have identified beta 1 and alpha 10 integrins, along with α-parvin, as physical regulators governing cell polarity and rotation during column formation (*30, 34, 36*). Furthermore, studies in embryonic chick and mouse limbs have shown that cell surface signaling through the Fz/Vangl/PCP pathway plays a major role in regulating chondrocyte polarity and rearrangement (*18, 19, 26, 28, 29, 37*) and that GDF5 is involved in chondrocyte orientation (*7*). Finally, studies in paralyzed mice (*38*) and muscle-less mouse embryos (*27, 39*) have uncovered the important role of muscle load in regulating cell polarity and column formation.

In this study, we analyze the 3D architecture of confetti-labeled clones in the embryonic and postnatal growth plate of mice. Intriguingly, we found that chondrocytes in the embryonic growth plate are not arranged in columns. Instead, confetti-positive clones give rise to elongated clusters oriented orthogonally to the P-D bone axis, as a result of non-stereotypic cell stacking. However, in the postnatal growth plate, confetti-positive clones form complex columns through a combination of stereotypical and non-stereotypic cell stacking, as well as small orthogonally oriented clusters. Additionally, we observed that column formation is buffered, permitting deviations of up to 60% misrotation between successive cells within columns. Our discovery that columns are absent embryonically implies that longitudinal bone growth is governed by different cellular mechanisms during embryonic and postnatal development, while highlighting the imperative role of 3D analysis when studying complex cellular arrangements such as columns.

## RESULTS

### 3D imaging of cell clones in the embryonic growth plate reveals stacking patterns that do not support column formation

To date, a comprehensive 3D analysis of column formation in the mouse embryonic growth plate has not been performed. To address this gap, we conducted multicolor clonal lineage tracing on proximal tibia and distal femur growth plates of *Col2a1-CreER:R26R-Confetti^+/-^* embryos.

Labeled cellular clones were subjected to 3D morphometric analysis using our previously reported 3D MAPs pipeline (*7*) (Supplementary Fig. 1A and 1B). Recombination was induced at E14.5 and four days later, chondrocyte clones were observed in all growth plate zones (Fig. 1A- C). 2D optical sections from both femur and tibia showed that cells within the clones were neatly stacked, forming a column-like structure that appeared parallel to the P-D bone axis (Fig. 1B,C). However, 3D examination revealed no columnar organization and cells that were rarely stacked neatly (Fig. 1D,E).

**Figure 1:**
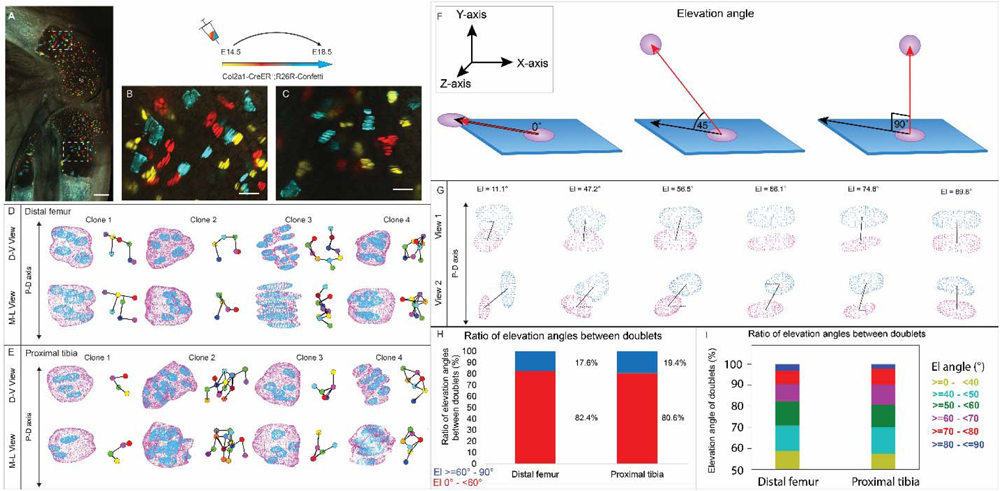
3D imaging of clones in the embryonic growth plate reveals complex morphologies. Chondrocyte clones in the proximal tibia and distal femur growth plates of *Col2a1-CreER^T2^:R26R-Confetti*^+/-^ mice were pulsed by tamoxifen administration at E14.5 and imaged at E18.5. **A-C** An image of chondrocyte clones in the knee was captured with a combination of multiphoton and confocal imaging using a Leica TCS SP8 confocal laser-scanning/MP microscope. Sparse labeling is observed throughout the growth plate. Scale bar: 250 µm. Magnified optical section of distal femur (**B**) and proximal tibia (**C**) highlight clones in the proliferative, prehypertrophic and hypertrophic zones, which appear to form columns. Scale bars: 50 µm. **D, E** 3D rendering of representative clones along the D-V and M-L axes from the distal femur (**D**) and proximal tibia (**E**) growth plates. Clone surface is in magenta and nuclear surface in blue. Skeletonized illustrations on the right highlight the complexity of clonal morphologies. Nuclear centroids are depicted as randomly colored circles; lines connect between nearest neighbor nuclei. **F** Illustration of various elevation angles between two cells. An elevation angle of 0° indicates that the two cells are in the same XZ plane, whereas an elevation angle of 90° indicates that the cells are in the same XY plane. **G** Representative images of nuclei at different elevation angles in two orthogonal viewing angles. Solid black lines represent the shortest distance between nuclear centroids. Elevation angle is the angle between the dashed black line and solid black line. **H, I** Stacked histograms show quantification of elevation angles between doublet cells in distal femur (n = 1044) and proximal tibia clones (n = 805). **H** Proportion of complete rotations (i.e., elevation angles of 60°-90°, in blue) vs partial rotations (under 60°, in red). **I** Distribution of elevation angles (°) is color-coded as indicated. Three biologically independent samples were examined in nine independent experiments.

**Table 1.**
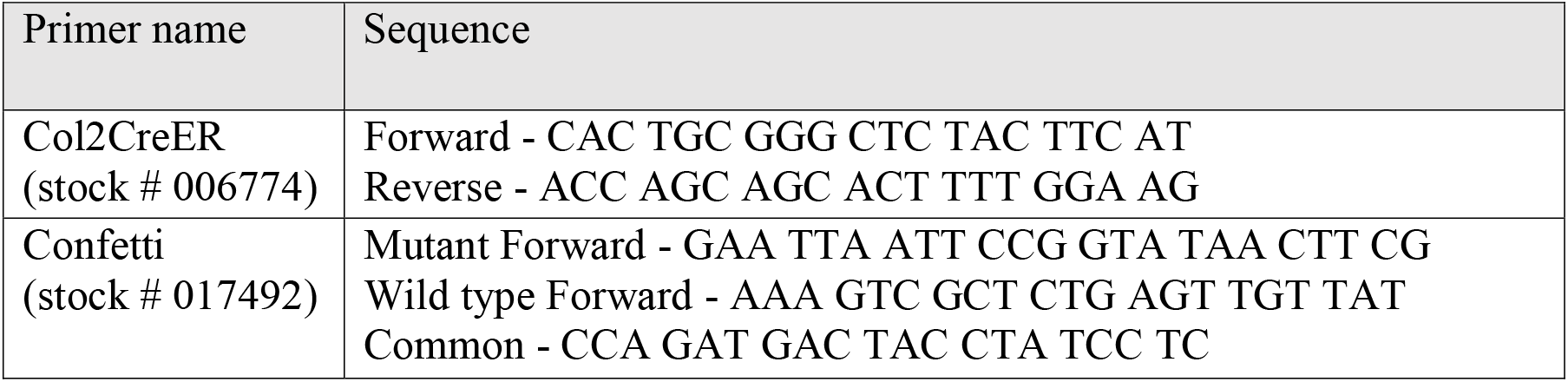
List of primers used for genotyping (all available from Jackson Laboratories).

To characterize the stacking behavior of embryonic growth plate cells, we performed quantitative analysis of local cell stacking in distinct clones. For that, we measured the elevation angle between pairs of cells (doublets) within a given clone (see details in Methods and Figure 1F). In a spherical coordinate system, the elevation angle between two perfectly stacked cells would be 90° (Figure 1F), indicating a complete rotation of the division plane during cell division. Visualization of cell doublets representative of various elevation angles employing two orthogonal viewing directions revealed that in 3D, typically stacked column cells are expected to exhibit elevation angles in the range of 60°-90° (Figure 1G). Notably, however, quantification of elevation angles revealed that less than 20% of doublets within a clone were typically stacked (distal femur, 17.6%; proximal tibia, 19.4%; Figure 1H,I). These results show that clone cells in the embryonic growth plate do not display the typical stacking behavior associated with a columnar arrangement, as most cells undergo partial division plane rotation.

### Columns are rare in the embryonic growth plate

Our finding that embryonic clones did not exhibit typical cell stacking characteristic of columns raised the question of their contribution to bone elongation. Atypically stacked cells, with elevation angles less than 60°, could still support longitudinal growth if the clones have elongated morphologies along the P-D axis of the growth plate. Thus, to characterize clone morphology, we extracted the long, medium, and short axes and measured the ratios between them (Fig. 2A; see Methods). Ratios close to 1 across all axes would indicate a spherical shape, whereas ratios approaching 0 would reveal a flattened ellipsoidal shape. Results showed that the long axis of the clones was consistently at least twice the length of the short axis. Moreover, in half of the clones the long axes measured at least five times the length of the short axes (distal femur, 58%; proximal tibia, 47%), indicating ellipsoid morphologies (Fig. 2C). Further examination showed that in roughly half of the clones, the long axis was at least twice the length of the medium axis (distal femur, 58.4%; proximal tibia, 54.3%; Fig. 2B) and the medium axis was at least twice the length of the short axis (distal femur, 58.5%; proximal tibia, 50.4%; Fig. 2D). Altogether, these results indicate that embryonic clones are either lentil-shaped oblate ellipsoids or rugby ball-like prolate ellipsoids and, thus, may contribute to bone elongation.

**Figure 2:**
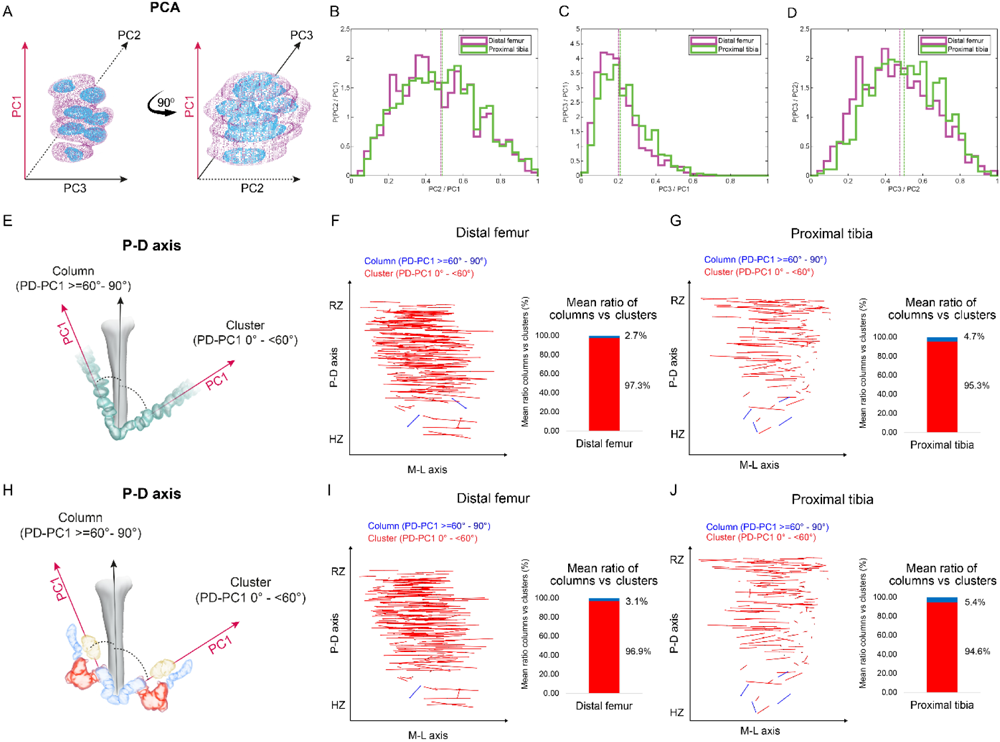
Columns are rare in the embryonic growth plate. Clone morphology was extracted by calculating the three orthogonal axes of each clone using PCA. PC1 (pink arrow) represents the long axis of the clone, PC2 (dashed black arrow) the medium axis, and PC3 (solid black arrow) the short axis. **A** A schematic drawing of the same clone from two orthogonal viewing angles with principal components labeled. **B-D** Histograms of clone PC ratios in E18.5 distal femur (DF) and proximal tibia (PT) growth plates reveal that clonal morphology is either oblate or prolate ellipsoid. In half of the clones, the long axis was at least twice the size of the medium axis (PC2/PC1: DF mean ± SD, 0.464 ± 0.197; PT, 0.469 ± 0.201; B), the long axis was at least five times the size of the short axis (PC3/PC1: DF, 0.201 ± 0.109; PT, 0.226 ± 0.119; C), and the medium axis was at least twice the size of the short axis (PC3/PC2: DF, 0.456 ± 0.189; PT, 0.504 ± 0.182; D). Dashed lines show the mean between samples. **E** Scheme illustrating the threshold between uniclones considered as columns (i.e., angle between long axis of the clone and P-D axis of the bone is 60°-90°) or clusters (i.e., angle is below 60°). **F, G** Orientation maps along the P-D and M-L axes and quantification of mean ratio of columns (in blue) vs clusters (in red) per sample for clones in the DF (n = 1044; **F**) and PT (n = 805; **G**) growth plates. Each line in the map represents the orientation of the long axis of an individual clone, whereas the length of the line is proportional to that of the clonal long axis (PC1). RZ refers to the middle of the resting zone; HZ refers to the end of the hypertrophic zone. **H** Scheme illustrating the threshold between multiclones considered as columns or clusters. **I, J** Orientation maps and quantification of mean ratio of multiclonal columns vs clusters per sample for the DF (n = 816; I) and PT (n = 619; J) growth plates.

Next, we sought to determine whether the elongated embryonic clones are aligned with the P-D bone axis. Previous studies suggested that in a 2D Cartesian coordinate system, single columns orient their long axis within 12° of the P-D bone axis, whereas multicolumns, i.e. those composed of multiple cell stacks, orient within 20° (*19*). In a 3D spherical coordinate system, these values correspond to elevation angles of 78° and 70°, respectively. We therefore set a more permissive threshold of 60° elevation to determine whether or not a clone qualifies as a column (Figure 2E). Measurements of the angle between the long axis of the clone and the P-D bone axis (see Methods) revealed that nearly all clones in the proximal tibia (mean, 95.4%) and the distal femur (mean, 97.3%) oriented perpendicular to the P-D axis (Figure 2F,G, and Supplementary Figure 2B-D). On average, only 4.6% of clones in the proximal tibia and 2.7% in the distal femur displayed a column-like orientation. (Figure 2F,G and Supplementary Figure 2C). Together, these results show that while embryonic clones have elongated morphologies, they do not support longitudinal growth.

### Multiclones do not form columns

Previous studies have shown that embryonic columns may be formed by merging of multiple clones (*19, 31*). This opens the possibility that in the embryo columns are multiclonal. To examine this possibility, we allowed neighboring clones to join (see Methods) and then performed orientation analysis, measuring the angle between the long axis of the multiclone with the P-D bone axis as before (Figure 2H). As depicted in Figure 2 and Supplementary Figure 3, nearly all multiclones in the proximal tibia (mean, 94.6%) and distal femur (mean, 96.8%) were oriented perpendicular to the P-D axis (Figure 2I,J, and Supplementary Figure 3A-D). On average, only 5.37% of multiclones in the proximal tibia and 3.12% in the distal femur were aligned parallel to the P-D bone axis, thereby satisfying the global orientation criterion for a column (Figure 2I,J). Together, these results show that embryonically multiclones do not form columns.

### 3D imaging of postnatal growth plate clones reveals diverse complex morphologies

Having found that embryonic clones do not meet the criteria for columns, we proceeded to analyze the 3D clonal structure in postnatal growth plates. For that, we used the same pipeline (*7*) to analyze proximal tibia and distal femur growth plates from clonally labeled *Col2a1- CreER:R26R-Confetti^+/-^* mice (Supplementary Figure 1A) at P40, 10 days after Cre induction. 2D optical sections showed small clones in the resting zone directly beneath the secondary ossification center, alongside longitudinal clones that spanned most of the growth plate height (Figure 3A-C). However, 3D rendering of confetti clones coupled with maps, generated by using multiple viewing angles, highlighted the diversity and complexity of these clones (Figure 3D,E). Whereas the small clones expanded along the D-V and M-L axes (Figure 3D), the longitudinal axis of most large clones visually aligned with the P-D bone axis. Surprisingly, however, nearly all the large clones, which contained 15 - 100 cells, had a complex morphology that was apparent from a particular viewing angle. Additionally, each large clone displayed motifs along its length, where cells appeared to stack typically before branching off into a horizontal expansion.

**Figure 3:**
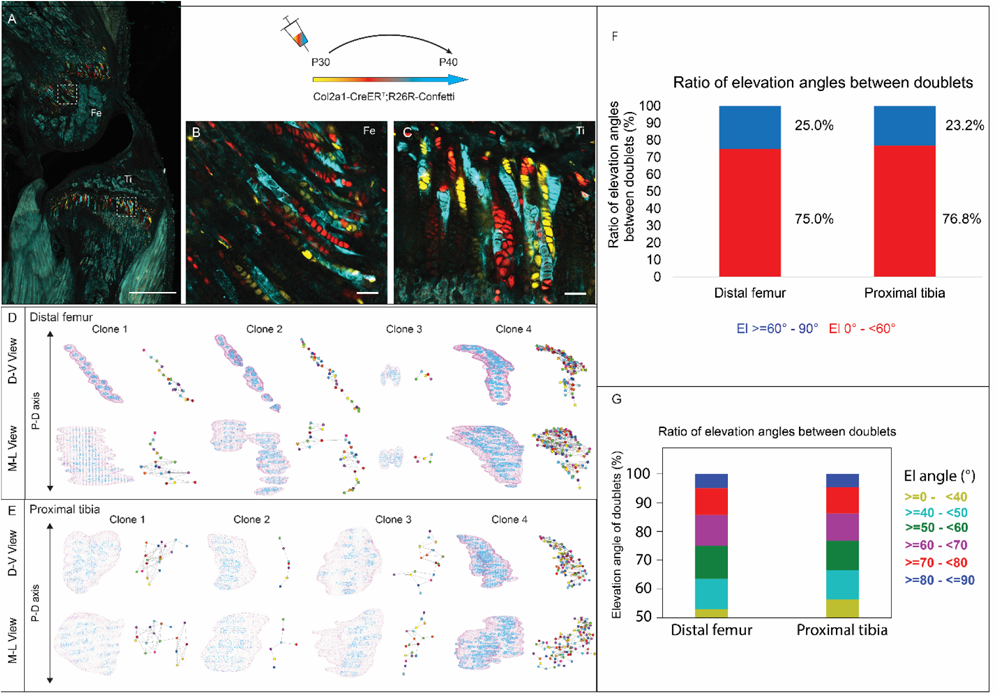
3D imaging of clones in the postnatal growth plate reveals diverse and complex morphologies. 3D morphology of chondrocyte clones was analyzed in the PT and DF growth plates of *Col2a1-CreER^T2^:R26R-Confetti* mice. Cells were pulsed by tamoxifen administration at P30 and traced until P40. **A** An image of chondrocyte clones in a P40 mouse knee was captured with a combination of multiphoton and confocal imaging using a Leica TCS SP8 confocal laser-scanning/MP microscope. Sparse labeling is observed throughout the growth plate. Scale bar: 1 mm. **B, C** Magnified optical sections of DF and PT clones reveal complex clones that appear to form columns. Scale bars: 50 µm. **D, E** 3D rendering of representative clones along the D-V and M-L axes of the DF and PT growth plates. Clone surface is in magenta and nuclear surfaces in blue. Skeletonized illustrations on the right highlight the complexity of each clone. Nuclear centroids are depicted as a randomly colored circle; lines represent connections with nearest neighbor nuclei. **F**, **G** Stacked histograms show quantification of elevation angles between cell doublets in clones. (F): Ratio between good rotations (El, 60°-90° in blue) and misrotations (El, 0°- 60°, in red). **G** Distribution of elevation angles (°), color-coded as indicated. Distal femur, 1866 clones; proximal tibia, 1666 clones. Three biologically independent samples were examined in nine independent experiments.

### Postnatal clones lack stereotypical cell stacking

Next, to determine whether postnatal clones form columns, we examined the two criteria for columns, namely local cell stacking and global orientation parallel to the P-D axis. To determine the degree of local cell stacking in the postnatal clones, we analyzed the elevation angle between all pairs of cells within a clone (Figure 3F,G). Results showed that less than 30% of doublets were typically stacked with elevation angles greater than 60° (distal femur, 25%; proximal tibia, 23.2%; Figure 3F). In addition, more than half of the doublets oriented orthogonally to the P-D axis (Figure 3G). These results are surprising given the observed elongated morphologies of clones. Interestingly, perfect rotations, characterized by elevation angles between 80°-90°, which were previously predicted to be prevalent (*19*), were rare (5.8% in the DF and 5.6% in the PT; Figure 3G).

### Complex longitudinal clones function as columns in the postnatal growth plate

Next, we studied the global orientation of postnatal clones by measuring the angle between the long clone axis and the P-D bone axis (Figure 4 and Supplementary Figure 4A,B). Interestingly, the results revealed the presence of clones with two different morphologies, i.e. columns that aligned to the P-D bone axis (PT: 30.5% and DF: 43.9%) and clusters, which oriented orthogonally (PT: 69.4% and DF: 56.1%) (Figure 4C,G and Supplementary figure 4C). To assess the possible functions of clusters and columns, we analyzed clone size. We found that columns varied in size, ranging from 2 to over 100 cells, many of which traversing the entire length of the growth plate (Figure 4D and 4H). By contrast, most clusters were composed of 2-10 cells and were located directly beneath the secondary ossification center in the RZ and at the very end of the HZ (Figure 4A,B,E,F and Supplementary figure 4A). While large columns likely contribute to longitudinal bone growth, the function of small clusters is unclear.

**Figure 4:**
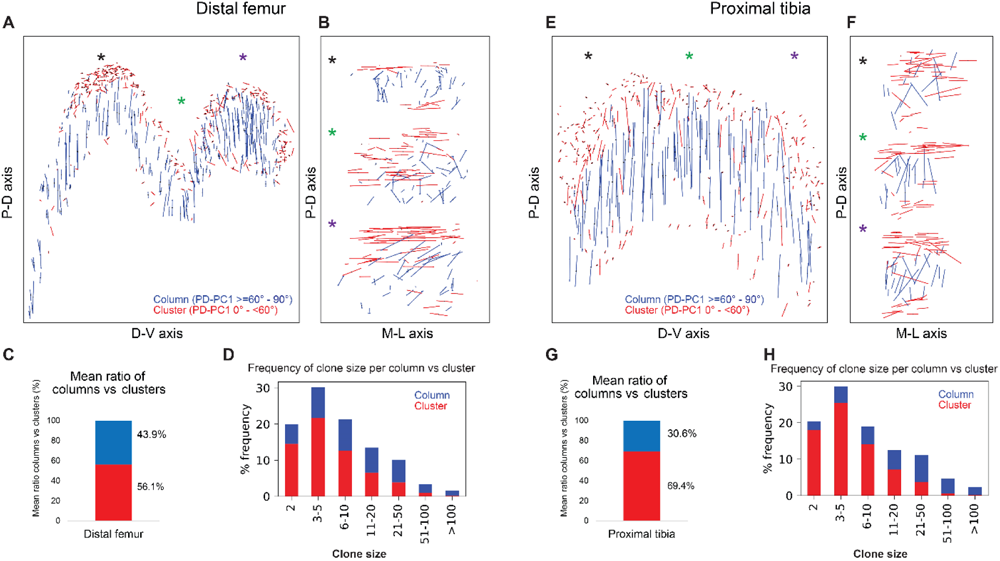
Complex longitudinal clones function as columns in the postnatal growth plate. Orientation maps of clones in P40 growth plates. **A, B** Clone orientation along the P-D and D-V axes of DF growth plates (n = 737 columns, 1129 clusters). Asterisks indicate the same locations in the growth plate. Each line represents the long axis of an individual clone, with its length proportional to that of the clone long axis. Columns are shown in blue and clusters in red. **C** Quantification of mean ratio of columns vs clusters in the DF growth plates. **D** Frequency of clone size per column (blue) vs cluster (red). **E, F** Clone orientation in the PT growth plates (n = 512 columns, 1154 clusters). **G**, **H** Mean ratio of columns vs clusters and frequency of clone size in PT growth plates. Three biologically independent samples were examined in nine independent experiments.

### A column can tolerate 60% misrotations

The main mechanism driving column formation is the rotation of the division plane between sister cells during oriented cell division (*19, 20, 30*). Our findings of two distinct morphologies of postnatal clones and non-stereotypic stacking patterns in embryonic growth plate clones raised the question of the rotational threshold that is required to maintain a columnar structure. To determine the rotation between pairs of cells, we assumed that the final orientation is dictated by the division plane rotation (*19, 20, 30*). Thus, a 90° division plane rotation would result in an elevation angle of 90°, whereas 0° would indicate no rotation (see Methods). As before, we classified elevation angles exceeding 60° as complete rotations, signifying a typically stacked cell doublet oriented along the P-D axis in at least two orthogonal viewing angles (Figure 1G). Analysis of hundreds of columns (distal femur, n = 737; proximal tibia, n = 512) revealed that rotations are successful roughly 40% of the divisions (distal femur, 39.5%; proximal tibia, 36.4%) (Figure 5A,C). Perfect rotations (80°-90°) occurred in less than 10% of the cases (distal femur, 9.6%; proximal tibia, 8.2%) (Figure 5B,D). By contrast, clusters exhibited only 15.5% complete rotations in the distal femur and 17.3% in the proximal tibia, with perfect rotations observed in 1.9% and 3% of divisions, respectively.

**Figure 5:**
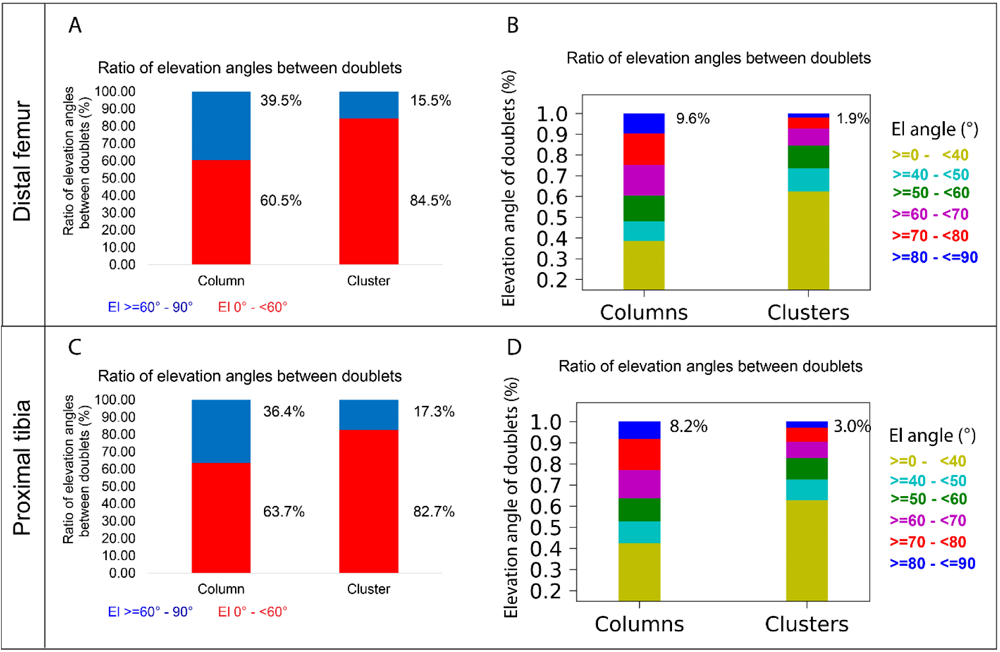
A column can tolerate 60% misrotations. **A, B** Stacked histograms show the proportion of doublet cells exhibiting complete rotations (i.e., elevation angle of 60°-90°, in blue) vs misrotations (El under 60°, in red; A) and distribution of elevation angles in columns vs clusters (B) in P40 DF growth plates (n = 737 columns, 1129 clusters). **C**, **D** Same analysis in PT growth plates (n = 512 columns, 1154 clusters). Elevation angles (°) are color-coded as indicated.

Lastly, we analyzed the elevation angle as a function of column or cluster size to explore the potential relationship between the two (Supplementary Figure 5). Although we did not observe a relationship between cluster size and misrotation, we found that the proportion of misrotations increases with column size. Altogether, these results suggest that column formation is resilient, capable of tolerating 60% misrotations. As misrotations accumulate in a clone with every cell division and, subsequently, cross this tolerance threshold, the structure will expand orthogonal to the P-D axis and form a cluster.

## DISCUSSION

Chondrocyte columns are a hallmark of growth plate architecture that, in turn, drives bone elongation. In this work, we studied column formation by analyzing the 3D structure of cell clones in the embryonic and postnatal growth plate of mice, using a modified version of 3D MAPs (*7*). Addressing the three criteria for a column, we found that uniclonal and multiclonal columns do not form in the embryonic growth plate. Instead, clones form elongated clusters that orient orthogonal to the P-D bone axis, as a result of numerous misrotations that occur during cell division within a clone. Postnatally, clones form complex columns built from a combination of ordered and disordered stacks of cells, as well as small orthogonally oriented clusters. We found that column formation is buffered, allowing at most 60% misrotations within columns (Figure 6).

**Figure 6:**
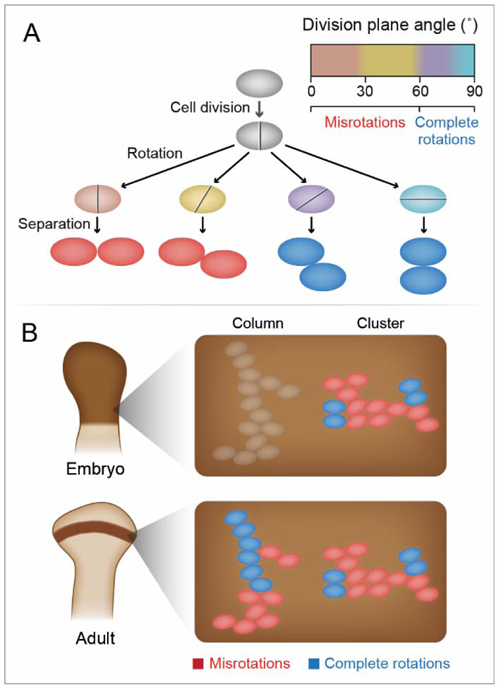
Model of column and cluster formation. **A** During oriented cell division in the growth plate, the division plane rotation can range from 0° to 90° (brown – blue). Less than 60° rotation indicates a misrotation (red), whereas rotations greater than 60° indicate a complete rotation (dark blue). **B** Columns form when 40% of division plane rotations are complete. Clusters form when 80% of division plane rotations are incomplete. In the embryonic growth plate, only clusters form, whereas in the postnatal growth plate both columns and clusters form.

Column formation is commonly viewed as a key morphogenetic process during bone elongation, in which clones of PZ cells stack with their short axis parallel to the P-D bone axis, resulting in the long axis of the column laying parallel along the P-D axis. This arrangement restricts lateral cell density, thereby maximizing the effect of chondrocyte hypertrophy along the P-D axis (*25*). Our discovery that during embryogenesis, when bone elongation is at its highest, the growth plates have no columns clearly requires a new thinking about the mechanisms of bone elongation. In the absence of columns, increase in cell volume is the most likely engine of longitudinal growth. Changes in chondrocyte morphology, proliferation, and matrix secretion may also play important roles in this process; however, their exact contributions to embryonic bone elongation are unclear.

Our finding that embryonic clones formed elongated clusters that oriented orthogonal to the P-D axis is interesting, as it may relate to the role of the growth plate in regulating morphological features like bone circumference and curvature. In contrast to bone elongation, the cellular mechanism by which the growth plate regulates these features are poorly studied. An example of the possible regulation of bone circumference by growth plate cells is that disruption of cell polarity often leads to wider bones (*7, 26, 28, 34, 40*). Another example is the finding that chondrocytes in growth plates of *Gdf5* mutant mice display abnormal cell orientation, which correlates to aberrant bone curvature (*7*).

Whereas in the embryonic growth plate clones did not form columns, postnatally we observed both clones organized in small orthogonally oriented clusters and large, complex columns containing a mixture of stereotypic and nonstereotypic cell stacks. The spatial segregation we observed between these two organizations suggests they may represent mechanical heterogeneity or serve different functions (*41*). Columns, which are located in the core, may contribute to bone elongation as suggested previously, whereas clusters may contribute to other morphogenetic functions of the growth plate. Interestingly, previous work on column formation in articular cartilage found clones were built from nonstereotypic cell stacking, raising the question of the function they may serve (*42*).

The core mechanism that underlies the column formation is the rotation of the division plane during cell division. Our findings that in the embryonic growth plate columns are not formed and that postnatal columns have complex morphologies along the medial-lateral and dorsal-ventral axes clearly suggests that 90° rotation of the division plane is uncommon. Indeed, our data show that in both embryonic and postnatal stages, only about one in ten divisions involved a full 90° rotation.

One limitation of our rotation analysis is that in order to calculate the relative position between neighboring cells, we assumed that chondrocyte rotation and orientation are correlated and that chondrocytes do not move after rotating, as was previously suggested (*20, 30*).

The finding from elevation angle analysis, that the mechanism that regulates the rotation of the division plane, which is the basis for column formation, works at less than 30% efficiency suggests two possibilities. The first is that this seemingly low efficiency is sufficient to allow proper growth plate activity. The other option is that inaccurate cell rotation fulfills a dual purpose during bone morphogenesis, promoting both longitudinal bone growth and other morphogenetic processes, such as circumferential bone growth.

The observation of chondrocyte columns in P40 growth plates raises the question of when columns start to form. Previous studies showed that there is a switch in column clonality when the secondary ossification center forms (*31–33*). Therefore, it is reasonable to speculate that this is also the time point at which clones switch from orthogonally oriented clusters to parallel columns.

Many of the studies on the cellular, molecular and mechanical mechanisms underlying column formation, e.g. Fz/Vangl/PCP signaling, integrins and muscle force, were conducted in embryos. In light of our discovery that columns do not exist at these stages, it will be interesting to revisit the role of these factors and pathways in the embryonic growth plate.

Here, we discovered that the core mechanism underlying chondrocyte column formation, namely a full rotation of the division plane during chondrocyte proliferation, is rare in the embryogenic growth plate. Because of partial rotations, columns do not form. Later, in the postnatal growth plate, the percentage of complete rotations of the division plane increases, resulting in the formation of complex columns in the center and small clusters in the periphery of the growth plate. Overall, these findings establish a new model for column formation and provide a new understanding of the cellular mechanisms underlying growth plate activity and bone elongation during development.

## METHODS

### Animals

For genetic labeling of chondrocyte clones in embryonic and postnatal growth plates, *Col2a1- CreER:R26R-Confetti^+/-^* mice were generate by crossing mice homozygous for *Col2-CreER^T^* (*43*) with R26R-*Confetti* homozygotes (Jackson Laboratories, B6.129P2- *Gt(ROSA)26Sor^tm1(CAG-Brainbow2.1)Cle^*/J; (*44*)). Mice were dissected in cold phosphate buffered saline (PBS), fixed for 3 hours at 4°C in 4% paraformaldehyde (PFA), washed in PBS and stored at 4°C in 0.5 M EDTA (pH 8.0, Avantor Performance Materials) with 0.01% sodium azide (Sigma) for 2 days. Limb samples were then dehydrated in 30% sucrose/PBS overnight at 4°C, embedded in OCT and stored at -80°C the following day. In all timed pregnancies, the plug date was defined as E0.5. For harvesting of embryos, timed-pregnant female mice were sacrificed by CO_2_ exposure. Embryos were sacrificed by decapitation with surgical scissors, and postnatal mice were sacrificed by CO_2_ exposure. Tail genomic DNA was used for genotyping by PCR (Supplementary Table 1). All animal experiments were pre-approved by and conducted according to the guidelines of the Institutional Animal Care and Use Committee (IACUC) of the Weizmann Institute. All animals used in this study had access to food and water ad libitum and were maintained under controlled humidity and temperature (45–65%, 22 ± 2°C, respectively). For each experiment, three mice were collected from at least two independent litters. Mouse embryos were used regardless of their sex, whereas postnatal experiments were performed only on females to control for potential sex-related phenotypes.

For clonal genetic tracing, tamoxifen was administered by oral gavage (Fine Science Tools) at a dose of 3 mg to P30 *Col2a1-CreER:R26R-Confetti^+/-^*mice or 2 mg to time-mated *R26R- Confetti* females at E14.5. Tamoxifen (Sigma-Aldrich, T-5648) was dissolved in corn oil (Sigma-Aldrich, C-8267) at a final concentration of 15 mg/ml. Neighboring cells that expressed the same fluorescent protein were considered clonal.

### Sample preparation

200-micrometer-thick sagittal cryosections of the embryonic or postnatal right hindlimbs from *Col2a1-CreER:R26R-Confetti^+/-^* mice were collected into a 12-well plate filled with 1 ml PBS. To remove OCT, samples were washed twice with PBS at room temperature (RT) with gentle rocking. Then, nuclei were stained with Draq5 (Thermo scientific 62252) diluted in PBST (PBS + 0.1% Triton X-100) for 2 hours at RT at a dilution of 1:2000 for embryonic samples and 1:1500 for postnatal samples. Three sections from the central region of the proximal tibia and distal femur growth plates were selected for further processing, together covering 600 µm along the medial-lateral bone axis. Sections were then placed in Rims (*7*) with a refractive index of 1.45 (74% Histodenz/PB) overnight at RT, and then mounted the following day with Rims between a glass slide and coverslip. Because Confetti fluorophores fade quickly, sections were imaged within one week of preparation.

### Image acquisition

The proximal tibia and distal femur growth plates were imaged by a combination of multiphoton and confocal imaging using an upright Leica TCS SP8 confocal laser-scanning/MP microscope (Leica Microsystems, Mannheim, Germany), equipped with external non-descanned detectors (NDD) HyD and HyD SP GaAsP detectors for confocal imaging (Leica microsystems CMS GmbH, Germany). Channels were collected in sequential mode. The CFP signal from the Confetti was excited by 900 nm laser line of a tunable femtosecond laser 680–1080 Coherent vision II (Coherent GmbH USA). Emission signal was collected using an external NDD HyD detector through a long pass filter of 440 nm. The GFP and YFP Confetti signal was excited by an Argon laser and collected with HyD SP GaAsP internal detectors (Ex 488 nm Em 498– 510 nm and Ex 514 nm Em 521-560 nm). The RFP Confetti signal was excited by a DPSS 561 nm laser with emission collection at 582-631 nm and the Draq5 signal was excited by a HeNe 633 laser, with emission collection at 669-800 nm. As reported previously (*31*), we rarely observed GFP clones in the growth plate.

Growth plates were imaged as a z-stack using a galvo scanner through a HC PL APO 20X/0.75 CS2 objective (scan speed, 400 Hz; zoom, 0.75; line average, 4; bit depth, 8; Z step, 0.39 µm). For embryonic samples, a format of 4096 x 4096 (XY) was used resulting in a pixel size of 180 nm (XY) and for postnatal samples, a format of 2000 x 2000 (XY) produced a pixel size of 369 nm (XY). Z stacks were acquired using the galvo scanner (objective movement) at 0.39 µm intervals.

Embryonic samples were imaged with a single field of view (FOV), which covered the middle of the resting zone through the beginning of the chondro-osseous junction (COJ). Postnatal samples were imaged with multiple overlapping FOVs (10% overlap), which covered the entire growth plate from the bottom of the secondary ossification center through the beginning of the COJ. Postnatal images were stitched in ImarisStitcher 9.9.0 (Bitplane).

### Growth Plate segmentation

To generate a mask of the growth plate, Microscopy Image Browser (version 2.81) (*45*) was used to manually segment the region between the secondary ossification center and the COJ in postnatal images and the entire growth plate region until the COJ in embryonic images.

Additionally, a mask of the hypertrophic zone was created by identifying the cells with the stereotypic chromatin staining unique to this zone.

### Nuclei segmentation

Images of fluorescently stained nuclei were automatically segmented as described previously (*7, 46*). For embryonic images a Gaussian blur filter (radius 2) and background subtraction (rolling ball radius 25) was applied in Fiji (*47*) prior to segmentation. We used standard deviations of σ = 12 for RZ, PZ and PHZ nuclei and of σ = 25 for HZ nuclei for embryonic images and σ = 5 for RZ, PZ and PHZ nuclei and of σ = 8 for HZ nuclei for postnatal images. Subsequently, local intensity maxima were extracted from the LoG-filtered image and reported as potential nuclear centers. For each potential seed point, we computed the mean intensity in a 4 x 4 x 4 voxel-wide cube for embryonic samples and 2 x 2 x 2 voxel-wide cube for postnatal samples surrounding the centroid. All image analyses were performed on a Windows Server 2012 R2 64-bit workstation with 2 Intel(R) Xeon(R)CPU E5-2687W v4 processors, 512 GB RAM, 24 cores and 48 logical processors.

### Clone segmentation

To generate a mask of each clone, .lif files were converted to .ims files using the Imaris file converter (version 9.8.0). Imaris surfaces tool was used to create surfaces and extract the three clone masks (yfp, rfp, and cfp) from the image; Gfp clones were not present in the images.

Surfaces were created with a grain size of 1.00 µm using the absolute intensity feature, with thresholds varying depending on the image. Surfaces with volumes greater than 150 µm^3^ were kept for further analysis. Next, the clone masks were overlapped with the raw signal in Fiji to inspect the quality of the segmentation. Quality was high in images where clones did not touch each other. In images with high labeling efficiency, where clones touched each other, some clone segmentations needed to be manually corrected in MIB.

### Nucleus and clone feature extraction

Following image segmentation and masking, separate segmented images were created for each clone mask in Fiji (S1 data) by assigning pixels outside of the clone mask and growth plate mask a value of 0. Next, segmented clone and nuclear images were relabeled using Morpholib plugin (*48*) and converted to 16- or 32-bit float depending on the number of objects in each image (S2 data). Images were reoriented in Fiji (*47*) so that the proximal-distal bone axis aligned to the Y-axis of the image coordinate. Nucleus and clone features were extracted as described previously (*7*) in Matlab (version R2017b) with a volume range of 100 – 1200 µm for nuclei and 150 – 500 x 10^7^ µm for clones.

### Morphometric analysis

Clone morphometrics, such as PC coefficients and PC orientations, were calculated as described previously (*7*). Cluster, column, or clone size was defined as the number of nuclei per cluster, column, or clone. These morphometric features were displayed as a histogram, a 3D morphology map, or both.

### Doublet quantification

In the clone analysis, the process starts by measuring distances between the centroids of all possible pairs of nuclei to identify the nearest neighbor pairs. These distances are sorted from low to high, establishing them as edges in the graph. For instance, in a clone with five nodes, there are 10 potential edges. The single-connected component method is applied to understand the graph’s topology and the distribution of nuclei within the clone. In a linear or columnar topology, a connected component should have a maximum of four edges (e.g., 1-2, 2-3, 3-4, 4- 5). In a spherical-like topology, a connected component can have all ten edges. The analysis begins with the smallest edge in the list. It is checked if this edge forms a single connected component. If not, the next smallest edge is added in the list, and this process continues. When a single connected component is achieved, the addition of edges stops. The final list of edges obtained in this manner defines the clone’s topology, with each edge termed as a doublet. These edges, present in the final list, are utilized as total number of doublets for the subsequent elevation angle analysis.

To evaluate the potential noise introduced into the measurement by measuring all nuclei pairs, the proportion of elevation angles as a function of number of nuclei neighbors per columns and clusters was calculated as well as the population statistics (Supplementary figure 6). We observed consistent patterns for both columns and clusters at any chosen number of nuclei neighbors. The range of the proportion of elevation angles between 60°-90° for 1-5 neighbors in columns was 13.2% - 15.5% for the distal femur and 16.3% - 21% for the proximal tibia. For clusters the range was 27.8% - 35.5% for the distal femur and 29.1% - 36.5% for the proximal tibia. The mean elevation angle distribution (60°-90°) from 1-10 neighbors is 14.5% (columns) and 28.4% (clusters) in the distal femur and 19.5% (columns) and 29.7% (clusters) in the proximal tibia (Supplementary figure 6A-D). The proportion of elevation angles up to 10 nuclei neighbors within columns and clusters represents up to 99.1% (Distal femur: 99.1% Proximal tibia: 98.8%) of all nuclei doublet pairs in columns and up to 71.6% (Distal femur: 71.6% Proximal tibia: 62%) in clusters (Supplementary figure 6E,6F). While examination of the influence of increasing neighbors on the proportion of elevation angles between 60°-90° did not show any statistical significance in columns, we found clusters to show statistical significance (p-value < 0.05, 2-sample t-test) when comparing the mean of lower number of neighbors (2–6) with larger numbers of nuclei neighbors (6–10) for both the distal femur and proximal tibia (Supplementary figure 6C,6D). This suggests that measuring elevation angle of nuclei doublets in clusters has slight noise present due to the spatial organization of nuclei in the clone, while the variation in columns is insignificant.

### Calculation of elevation angle and the angle between clone and PD axis

To determine the elevation angle between nuclei in a doublet, we first transformed the Cartesian coordinates of doublet to a spherical coordinate system. Then, we shifted the mean position of a doublet to the origin zero and used the MATLAB function cart2sph to obtain the elevation angle, azimuthal angle, and radius (r). If one nucleus in the doublet has an elevation angle of phi, the other nucleus’s angle is -phi. Similarly, if one nucleus has a positive azimuthal angle theta, the other has (theta-180). The elevation angle was measured from the +ve Z-axis towards the XY-plane and its values are in the range of [-pi/2, pi/2]. The azimuthal angle was measured in XY-plane from the +ve X-axis towards the +ve Y-axis, and its values are in the range of [-pi, pi]. The radius in the spherical coordinate system is equivalent to half of the distance between two nuclei in a Cartesian coordinate system. The elevation angle between two nuclei lying on top of one another is 90°, whereas the angle between two nuclei that are positioned next to each other in the XY plane is 0. Because the elevation angles of nuclei in a doublet mirror each other, we only used the absolute value.

To determine the distribution of elevation angles, we divided them into five categories: between 0° and <40° (color-coded in yellow), between 40° and <50° (cyan), between 50° and <60° (green), between 60° and <70° (magenta), between 70° and <80° (red), and between 80° and 90° (blue). The mean value of each category is reported as the elevation angle for column or cluster clone.

The angle between clone and PD axis (theta) was calculate using the MATLAB function “theta=atan2(norm(cross(u,v)),dot(u,v))”, where u and v are two vector representing PD axis and clone orientation. If the function value exceeded 90°, the reported value was (180°-theta). Finally, to use the same notation as for the elevation angle analysis, the PD-PC angle is reported as (90°-theta). Thus, if the longest axis of the clone (PC1) is parallel to PD axis, then the reported value is 90°, and if it is perpendicular, then the reported value is 0°.

### Analysis of elevation angle consistency

To evaluate the consistency of elevation angle computations between cells and nuclei, we used a dataset of a wild-type ulna sample (49) from our previous publication (7). This dataset included segmentation data on both nuclei and cells from the entire growth plate, but not clone information. We matched between cells and nuclei in the dataset by calculating pairwise distances between centroids and the angle differences along PC1 of cells and nuclei. Pairs were considered matched if the centroid distance was less than 10 micrometers and the PC1 angle difference was less than 45°, resulting in 3764 paired cells and nuclei.

To simulate random clones in the ulna sample, we generated clones of various sizes, radii, and random positions along the P-D axis. These parameters were sampled from real clone sizes in the embryonic distal femur sample. Within the pool of matched cell-nucleus pairs, we determined the number of cell centroid positions falling within the randomly sampled sphere radius and P-D positions, allowing a variation of up to 15% of the P-D height on either side. While cells meeting these criteria may or may not match the desired clone size, we calculated pairwise distances for these cells, sorting them from largest to smallest. Using the single-connected component method, detailed in the “Doublet quantification” subsection, we removed the farthest-apart cells and continued this process to obtain single connected components of cells with the desired clone size.

A total of 1143 random clones were generated in the paired data. Within these clones, we calculated the elevation angle for cell doublets and nuclei doublets separately and visualized the variation between them in a histogram (Supplementary figure 7A). The results indicated a striking similarity in histogram distribution. Additionally, we computed the Pearson correlation between the average elevation in a clone based on cell centroids and the average elevation in a clone based on nuclear centroids, resulting in a correlation value of 0.82 (Supplementary figure 7B). These results suggest that elevation angle calculation performed on nuclei can serve as a reliable proxy for cell-based measurements.

### Multiclone formation analysis through clone merging

In our analysis of embryonic data, clones were merged based on the criterion that the distance between any nucleus within one clone to any neighboring clone was less than 15 micrometers. After the merging process, the total number of nuclei in a merged clone equaled the sum of nuclei in the original individual clones. All properties of merged clones were computed in the same manner as for individual clones.

## DATA AVAILABILITY

The datasets generated and analyzed during the current study are available from the corresponding author on reasonable request.

## ACKNOWLEDGEMENTS

We thank Nitzan Konstantin for editorial assistance and members of the Zelzer lab for their advice and encouragement throughout this project. We thank Phillip T. Newton and Andrei S. Chagin for sharing Confetti labeled images for us to compare our data to. We thank M. Vijay Kumar, Uri Alon, Efi Efrati, Ariel Amir, Hillel Aharoni, and Jure Leskovec for useful discussions. We thank the de Picciotto Cancer Cell Observatory in memory of Wolfgang and Ruth Lesser, Weizmann Institute of Science, for providing confocal/multiphoton imaging infrastructure, Ishai Sher and Hanna Vega from the Graphic Design Department at the Weizmann Institute of Science for their help with graphics. This study was supported by grants from Israel Science Foundation (ISF) 1462/20, Breakthrough Research Grants (“MAPATZ”) 1387/23, Weizmann - Sagol Institute for Longevity Research, Weizmann - Kekst Family Institute for Medical Genetics, Weizmann – Belle S. and Irving E. Meller Center for the Biology of Aging, National Institutes of Health (NIH), UM/Israel Research Partnership, The David and Fela Shapell Family Center for Genetic Disorders Research, the Julie and Eric Borman Family Research Funds, the Yves Cotrel Foundation, and by the Nella and Leon Benoziyo Center for Neurological Diseases at the Weizmann Institute of Science.

## CONTRIBUTIONS

S.R. designed and carried out experiments, preprocessed the data and wrote the manuscript; A.A. designed and ran analyses and wrote the manuscript; A.S. carried out experiments and preprocessed the data; A.B. conceptualized analyses; P.V. conceptualized analyses; E.Z. designed and supervised experiments and analyses and wrote the manuscript. All authors reviewed the manuscript.

## ETHICS DECLARATIONS

The authors declare no competing interests.

