## Supplementary figures for "Bone elongation in the embryo occurs without column formation in the growth plate" (from another version of this preprint: version 1, 2023-11-16)

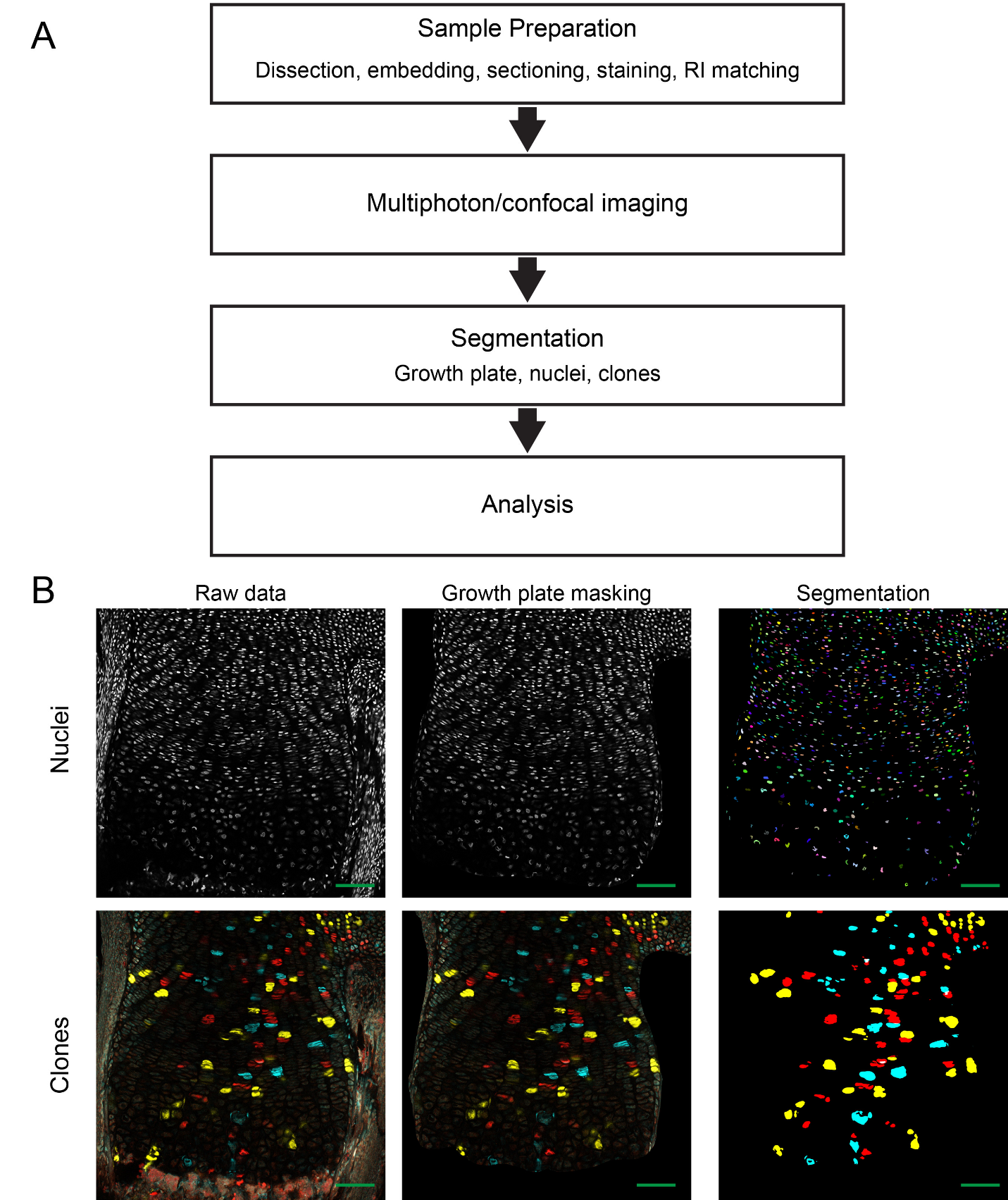

**Supplementary Figure 1: A** Scheme highlighting the different stages of the modified 3D MAPs pipeline. **B** For semi-automatic segmentation of clones and nuclei, the raw 3D image is masked in Microscopy Image Browser (MIB) to remove clones and nuclei not belonging to the growth plate. Then, the masked nucleus image undergoes automatic segmentation in XPIWIT to produce a randomly color-coded 3D image, where each color represents a single nucleus. The masked clone image undergoes semi-automatic segmentation in Imaris and Fiji, where each clone is colored according to its Confetti fluorophore (yellow, YFP; red, RFP; cyan, CFP). Scale bars: 100 µm.

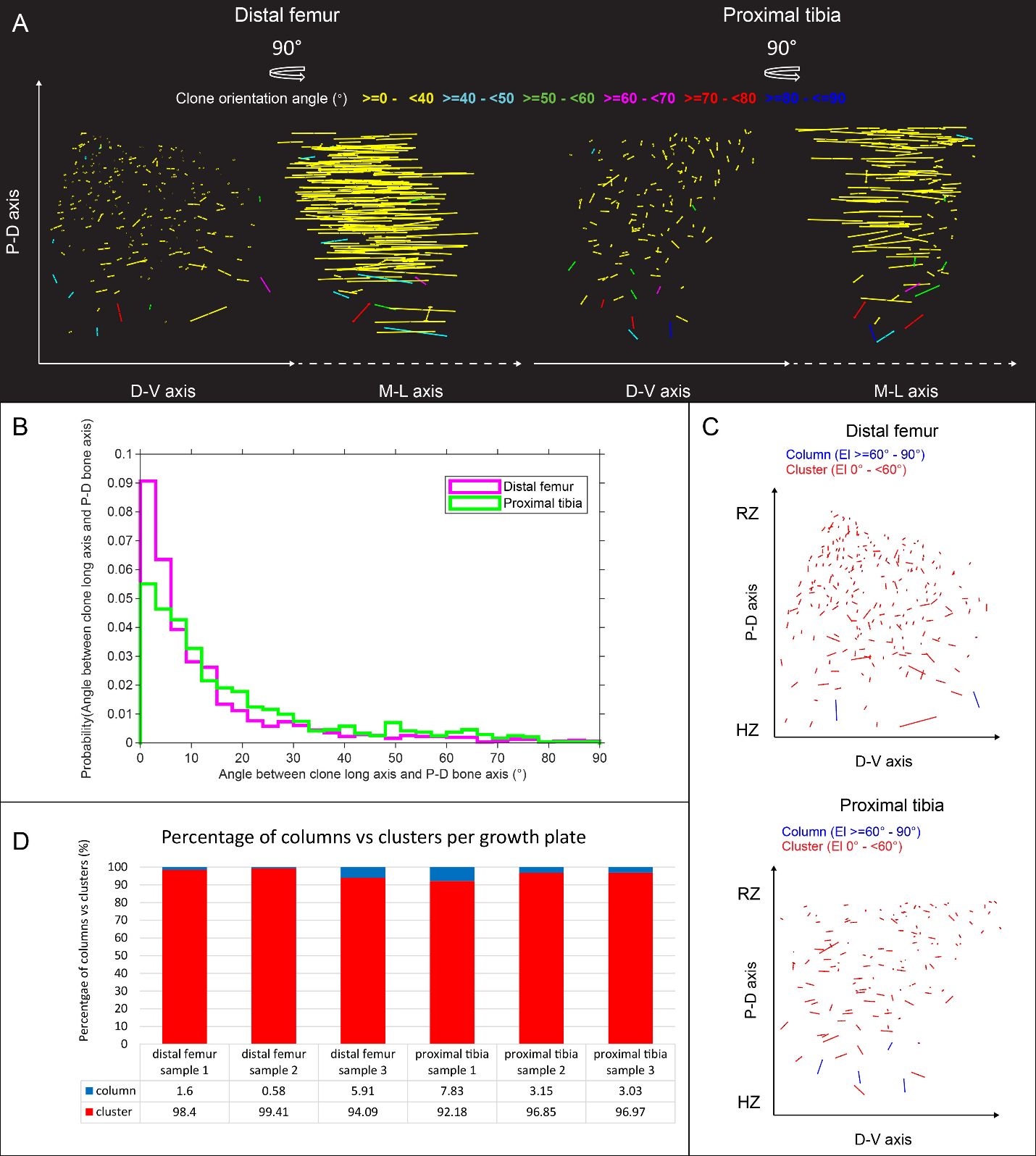

**Supplementary Figure 2: A** Orientation maps of uniclones from E18.5 DF and PT growth plates from two orthogonal viewing angles**.** Each line in the map represents the orientation of the long axis of an individual clone, whereas line length is proportional to that of the clone long axis (PC1). Clones whose long axis is parallel to the P-D bone axis are in dark blue, whereas and represents orthogonal clones are in yellow. Elevation angles (˚) are color-coded as indicated. **B** Graph showing distribution of elevation angles between the long axis of uniclones and P-D bone axis. **C** Orientation maps along the P-D and D-V axes of uniclonal columns (in blue) and clusters (in red) in the DF and PT growth plates. Each line represents the long axis of an individual uniclone, with its length proportional to that of the clonal long axis (PC1). RZ refers to the middles of the resting zone. HZ refers to the end of the hypertrophic zone. **D** Quantification of the percentage of uniclonal columns vs clusters per sample in the DF (n = 1044 clones) and PT (n = 805 clones) growth plates.

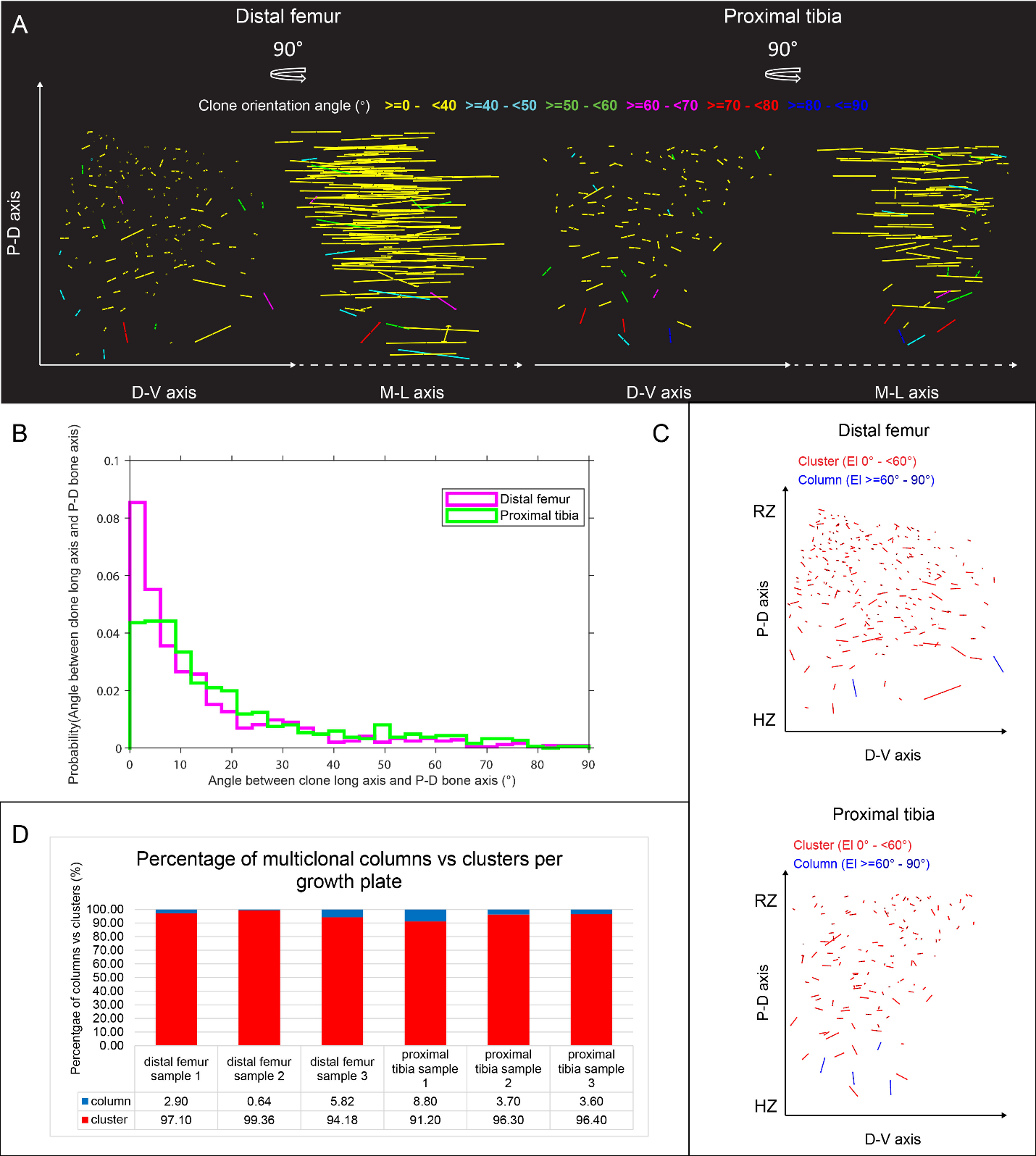

**Supplementary Figure 3: A** Orientation maps of multiclones in E18.5 DF and PT growth plates from two orthogonal viewing angles**.** Each line represents the long axis of an individual clone, with its length proportional to that of the clone long axis (PC1). Multiclones whose long axis is parallel to the P-D bone axis are in dark blue, whereas orthogonal multiclones are in yellow. Elevation angles (˚) are color-coded as indicated. **B** Distribution of elevation angles between multiclone long axis and P-D bone axis. **C** Orientation maps along the P-D and D-V axes of multiclonal columns and clusters in the DF and PT growth plates**.** Each line represents the long axis of an individual multiclone, with its length proportional to that of the clone long axis (PC1). Multiclonal columns (i.e., angle between clonal long axis and P-D bone axis is 60˚ - 90˚) are in blue, whereas multiclonal clusters (i.e. angle below 60˚) are in red. RZ refers to the middles of the resting zone. HZ refers to the end of the hypertrophic zone. **D** Quantification of the percentage of multiclonal columns vs clusters per sample in DF **(**n = 816 clones) and PT (n = 619 clones) growth plates.

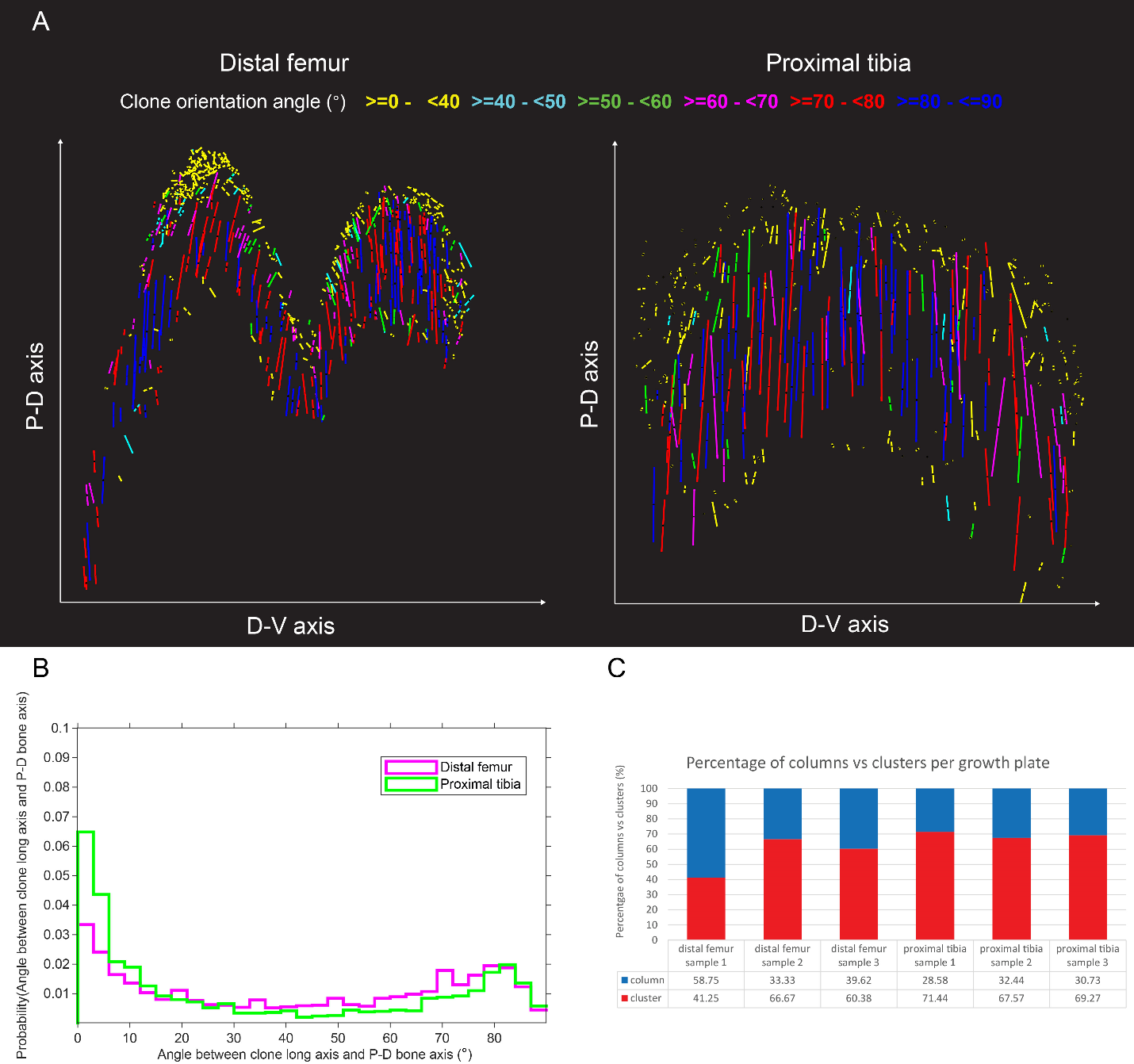

**Supplementary Figure 4: Clones orientation in the postnatal growth plate. A** Orientation maps along the P-D and D-V axes of single clones from DF (n = 1866) and PT (n = 1666) growth plates at P40**.** Each line represents the long axis of an individual clone, with its length proportional to that of the clone long axis. Clones that are parallel to the P-D axis are in dark blue, whereas orthogonal clones are in yellow. Elevation angles (˚) are color-coded as indicated. **B** Distribution of elevation angles between the clone long axis and P-D bone axis. **C** Quantification of the percentage of columns vs clusters per sample in DF and PT growth plates.

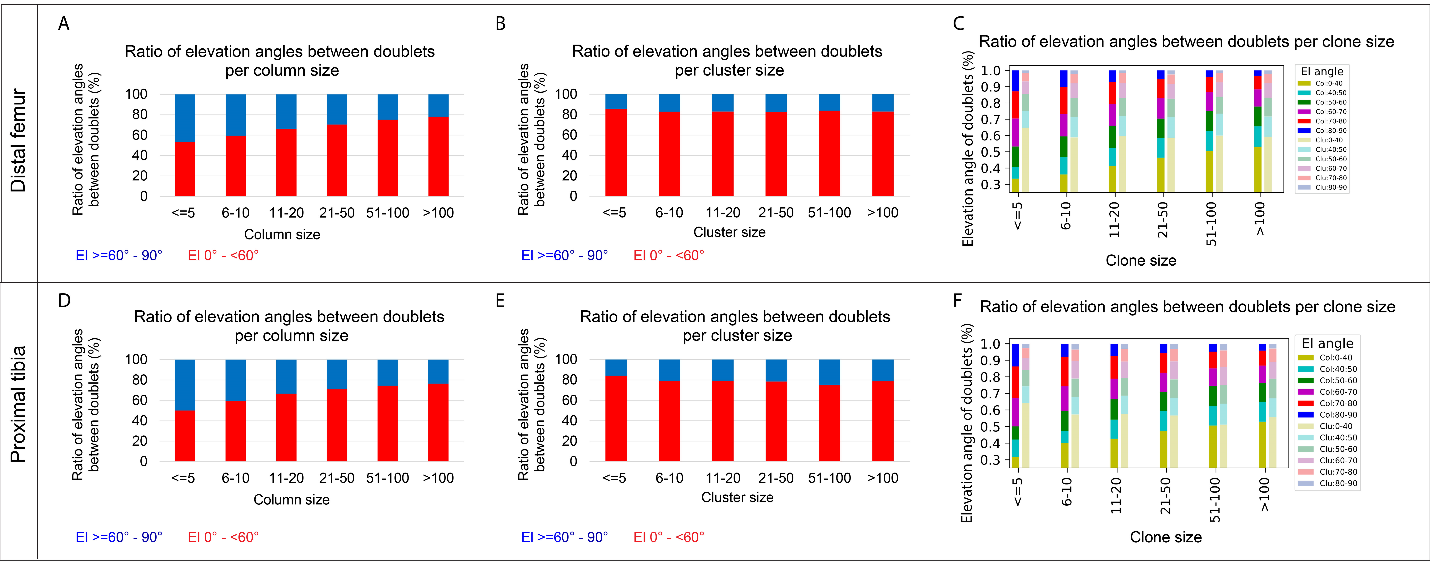

**Supplementary Figure 5. Elevation angles between postnatal doublet cells as a function of clone size.** **A-C** Quantification of elevation angles in cell doublets from P40 DF growth plates (n = 737 columns, 1129 clusters). **A, B** Stacked histogram shows the proportion of doublet cells exhibiting good rotations (i.e., elevation angle of 60˚- 90˚, in blue) vs misrotations (El under 60˚, in red) in columns (A) and clusters (B) as a function of column or cluster size. **C** Distribution of elevation angles between doublet cells as a function of clone size. **D-F** Analysis of cell doublets in P40 PT growth plates (n = 512 columns, 1154 clusters). Elevation angles (˚) are color-coded as indicated.

­­­­
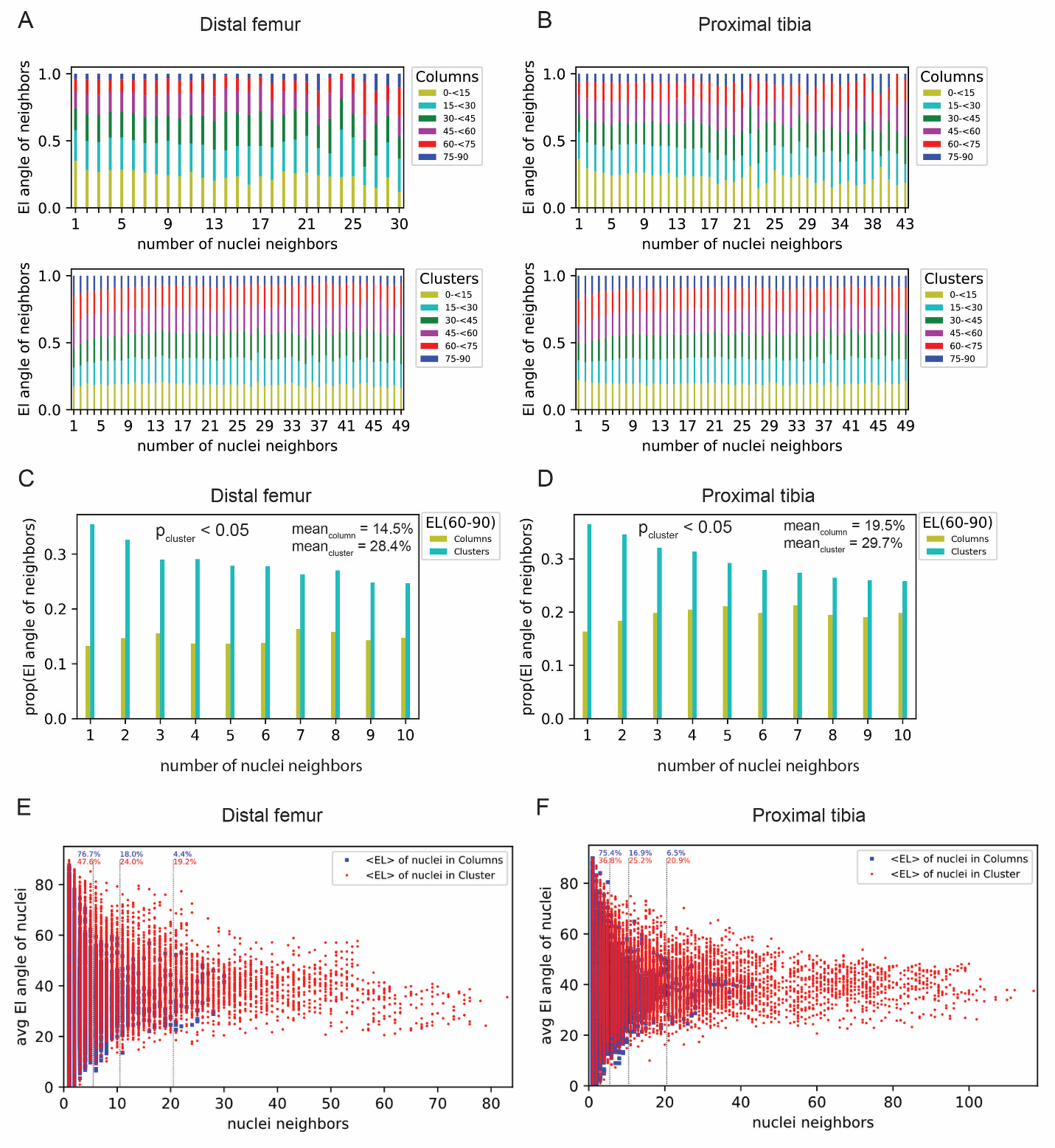

**Supplementary Figure 6. Noise evaluation of doublet analysis.** **A,B** Histogram showing the proportion of elevation angles from 0º-90º of nuclei per column and per cluster as a function of number of nuclei neighbors. **C,D** Histogram showing the proportion of elevation angles from 60º-90º of nuclei per column and per cluster as a function of number of nuclei neighbors. While examination of the influence of increasing nuclei neighbors on the proportion of elevation angles between 60º-90º did not show any statistical significance in columns, we found clusters to show statistical significance (p-value < 0.05, 2-sample t-test) when comparing the mean of lower number of neighbors (2-6) with larger numbers of nuclei neighbors (6-10) for both the distal femur and proximal tibia. DF, Mean_column_ = 14.5% Mean_cluster_ = 28.4%; PT, Mean_column_ = 19.5% Mean_cluster_ = 29.7%. **E,F** Histogram showing the distribution of mean elevation angle as a function of number of nuclei neighbors. Nuclei within columns in blue. Nuclei within clusters in red. DF column nuclei (n = 4872), DF cluster nuclei (n = 19121), PT column nuclei (n = 6856), PT cluster nuclei (n =18967). Dashed lines indicate the percentage of the population which falls under nuclei neighbor cutoffs of <=5, <=10, and <=20 neighbors respectively. El; elevation angle.

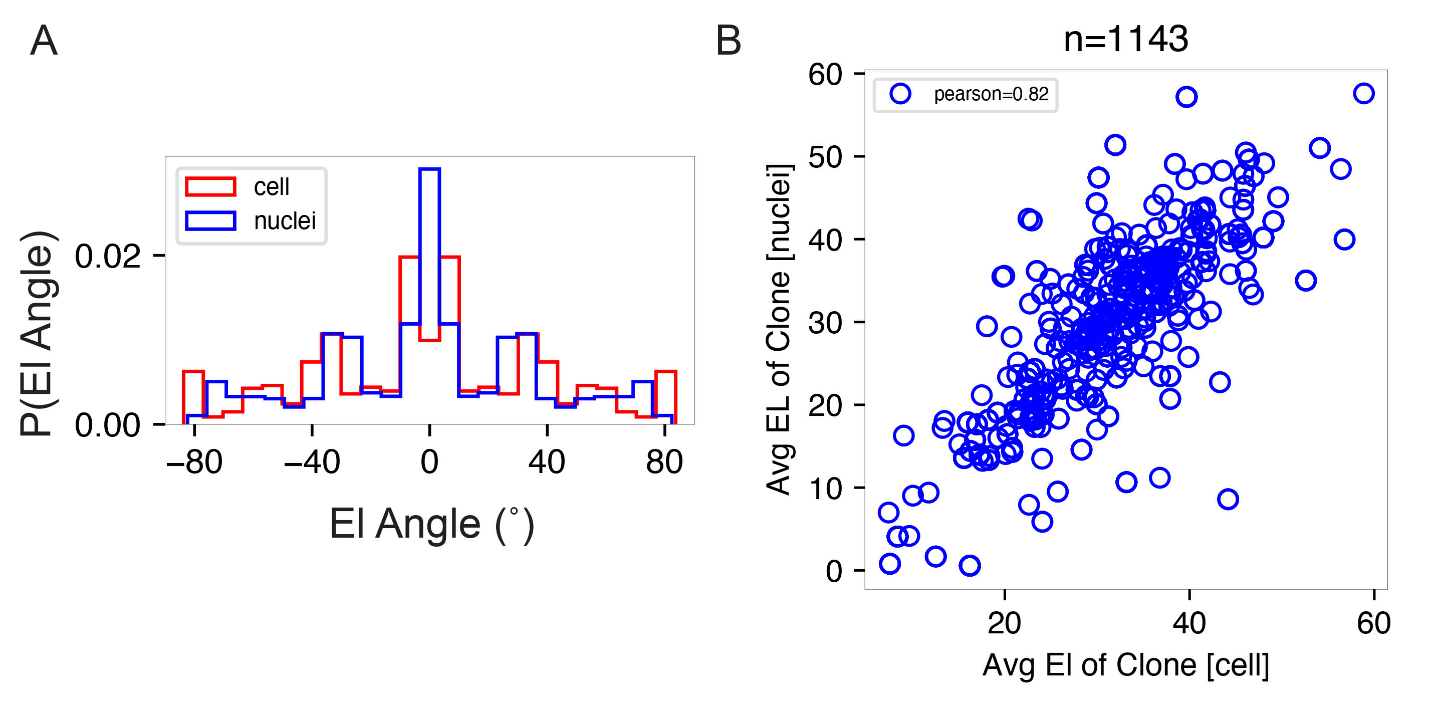

**Supplementary Figure 7. Correlation analysis between cells and nuclei.** **A** Histogram showing the distribution of elevation angles between doublet pairs of cells vs nuclei within a clone is nearly identical. Cells in red. Nuclei in blue. n = 1143 clones. **B** Pearson correlation analysis between cells and nuclei for elevation angle analysis. A Pearson coefficient of 0.82 suggests the nuclei can be used as a proxy for the cells. n = 1143 clones. El; elevation angle.
